# Unraveling the antiviral activity of plitidepsin by subcellular and morphological analysis

**DOI:** 10.1101/2021.12.16.472880

**Authors:** Martin Sachse, Raquel Tenorio, Isabel Fernández de Castro, Jordana Muñoz-Basagoiti, Daniel Perez-Zsolt, Dàlia Raïch-Regué, Jordi Rodon, Alejandro Losada, Pablo Avilés, Carmen Cuevas, Roger Paredes, Joaquim Segalés, Bonaventura Clotet, Júlia Vergara-Alert, Nuria Izquierdo-Useros, Cristina Risco

## Abstract

The pandemic caused by the new coronavirus SARS-CoV-2 has made evident the need for broad-spectrum, efficient antiviral treatments to combat emerging and re-emerging viruses. Plitidepsin is an antitumor agent of marine origin that has also shown a potent pre-clinical efficacy against SARS-CoV-2. Plitidepsin targets the host protein eEF1A (eukaryotic translation factor 1 alpha 1) and affects viral infection at an early, post-entry step. Because electron microscopy is a valuable tool to study virus-cell interactions and the mechanism of action of antiviral drugs, in this work we have used transmission electron microscopy (TEM) to evaluate the effects of plitidepsin in SARS-CoV-2 infection in cultured Vero E6 cells 24 and 48h post-infection. In the absence of plitidepsin, TEM morphological analysis showed double-membrane vesicles (DMVs), organelles that support coronavirus genome replication, single-membrane vesicles with viral particles, large vacuoles with groups of viruses and numerous extracellular virions attached to the plasma membrane. When treated with plitidepsin, no viral structures were found in SARS-CoV-2-infected Vero E6 cells. Immunogold detection of SARS-CoV-2 nucleocapsid (N) protein and double-stranded RNA (dsRNA) provided clear signals in cells infected in the absence of plitidepsin, but complete absence in cells infected and treated with plitidepsin. The present study shows that plitidepsin completely blocks the biogenesis of viral replication organelles and the morphogenesis of virus progeny. Electron microscopy morphological analysis coupled to immunogold labeling of SARS-CoV-2 products offers a unique approach to understand how antivirals such as plitidepsin work.

## INTRODUCTION

Infection caused by the severe acute respiratory syndrome coronavirus 2 (SARS-CoV-2) urgently needs effective antiviral treatments with a significant clinical benefit for hospitalized patients. So far, randomized clinical trials have failed to identify potent antivirals targeting the virus, with the only exception of remdesivir and molnupiravir, which have recently shown clinical benefits when administered early upon infection (Beigel et al., 2020; Garibaldi et al., 2021; Grein et al., 2020; Imran et al., 2021). As it happens with all viruses, coronaviruses have a reduced number of molecular druggable targets, and as new variants arise, these targets evolve and could eventually develop antiviral resistance. An interesting approach to overcome these limitations relies on the use of compounds against highly conserved cellular host factors required to complete the replication cycle of distinct types of viruses, which offer a common targeted solution to diverse viral threats. Furthermore, targeting host factors could offer a pan-antiviral strategy to combat not only viruses known at present, but also future pandemics to come (Baggen et al., 2021).

Presently, there are only a limited number of approved drugs involved in targeting host factors at post-entry steps (Baggen et al., 2021). This approach is especially relevant for pan-antiviral solutions given that viruses may use alternative pathways to enter a cell, but will most likely converge at intracellular processes involving genome replication and protein production. One of these compounds is plitidiepsin, which has shown a potent preclinical efficacy against SARS-CoV-2 by targeting the host protein eEF1A (Losada et al., 2016; Rodon et al., 2021; White et al., 2021). In 2018, the Therapeutic Goods Administration (TGA; Australian Government) approved the combination of plitidepsin with dexamethasone for the treatment of patients with relapsed/refractory multiple myeloma (https://www.tga.gov.au/auspar-plitidepsin). Currently, plitidepsin is being evaluated in a phase 3, multicenter, randomized, controlled trial to determine the efficacy and safety of two dose levels of plitidepsin *versus* control in adult patient requiring hospitalization for management of moderate Coronavirus infectious disease 2019 (COVID-19) (ClinicalTrials.gov Identifier: NCT04784559). eEF1A2 is necessary to transport aminoacyl-tRNAs to the A site of the ribosome during protein translation, but is also implicated in other activities (Mateyak and Kinzy, 2010) such as inhibition of apoptosis (Sun et al., 2014), proteasome degradation (Hotokezaka et al., 2002), and actin bundling and cytoskeleton reorganization (Edmonds et al., 1998) among other non-canonical functions. Also, eEF1A2 is implicated in the replication of distinct viruses, including coronaviruses (Zhang et al., 2014), and was identified as a potential SARS-CoV-2 interacting protein in one of the first screenings performed to identify novel targets (Gordon et al., 2020).

Upon SARS-CoV-2 infection, plitidepsin inhibits nucleocapsid viral protein expression and viral induced cytopathic effect *in vitro* (Rodon et al., 2021; White et al., 2021). In addition, it also reduces genomic and subgenomic RNA expression (White et al., 2021). Current models of SARS-CoV-2 replication propose that upon viral fusion, non-structural viral proteins form a replication– transcription complex that is continuous with the ER and has a double membrane vesicle (DMV) morphology that shelters the viral genome replication (Baggen et al., 2021; Wolff et al., 2020a). A negative RNA strand is used as a template for the generation of positive strands that are translated and incorporated into nascent viruses (Baggen et al., 2021; Wolff et al., 2020a). Discontinued transcription of positive RNA strands produce negative subgenomic RNAs, which are then used as templates for positive subgenomic RNA generation that codify for structural and accessory proteins (Baggen et al., 2021; Wolff et al., 2020a). Translation of viral proteins is facilitated by mRNA export via molecular pores located in the DMV that enable viral protein production in the cytoplasm (Wolff et al., 2020b). Yet, how plitidepsin exerts its intracellular antiviral activity and influences the formation of viral replication DMV remains unknown.

Here we aimed to explore the antiviral effect of plitidepsin at the cellular level to understand its impact on SARS-CoV-2 replication and DMV formation. Using transmission electron microscopy (TEM), we recapitulated the infectious SARS-CoV-2 cycle in Vero E6 cells and observed a lack of viral DMVs in plitidepsin-treated cells. Complementary immunohistochemistry analyses using nucleocapsid and dsRNA immunogold labeling unambiguously confirmed the lack of viral replication in plitidepsin treated cells.

## MATERIAL & METHODS

### Biosafety Approval

The biologic biosafety committee of the Research Institute Germans Trias i Pujol approved the execution of SARS-CoV-2 experiments at the BSL3 laboratory of the Center for Bioimaging and Comparative Medicine, CMCiB (CSB-20-015-M3).

### Materials

Plitidepsin was synthesized at PharmaMar, S.A. (Colmenar Viejo, Madrid, Spain).

### Cell culture, viral isolation and titration

Vero E6 cells (ATCC CRL-1586) were cultured in Dulbecco’s modified Eagle medium (Invitrogen) supplemented with 10% fetal bovine serum (FBS; Invitrogen), 100 U/mL penicillin, 100 μg/mL streptomycin (all from Invitrogen). SARS-CoV-2 D614G was isolated from a nasopharyngeal swab collected in March 2020 in Spain in Vero E6 cells as previously described in detail (Rodon et al., 2021). The virus was propagated for two passages and a virus stock was prepared collecting the supernatant from Vero E6 cell and sequenced as described elsewhere (Rodon et al., 2021). Genomic sequence was deposited at GISAID repository (http://gisaid.org) with accession ID EPI_ISL_510689. Viral stock was titrated in 10-fold serial dilutions on Vero E6 cells to calculate the TCID_50_ per mL.

### Cellular infection

Vero E6 cells were infected with SARS-CoV-2 at a multiplicity of infection (MOI) of 0.02 plaque forming units (PFU) per cell for 24 and 48 hours (h) in the presence or absence of 0.2 and 0.05 μM of plitidepsin added at the time of infection. These two concentrations are close to the IC_90_ and IC_50_ of plitidepsin, respectively, determined by the SARS-CoV-2 induced cytopathic effect on Vero E6 cells (Rodon et al., 2021). As negative controls, Vero E6 cells were treated with or without plitidepsin for the same time but without virus. At each studied time point, cell monolayers were chemically fixed using two different protocols depending on the type of analysis (morphology or immunohistochemistry).

### Morphology Analysis

Cell monolayers were fixed with 4% paraformaldehyde and 1% glutaraldehyde in phosphate buffered saline (PBS) for 2h at room temperature (RT). Cells were removed from the plates in the fixative, pelleted by centrifugation and washed three times with

PBS. Post-fixation of cell pellets was done on ice with 1% osmiumtetroxide + 0.8% potassium ferrocyanide in water. Afterwards the pellets were dehydrated on ice with increasing concentrations of acetone and processed for embedding in the epoxy resin EML-812 (Taab Laboratories), as previously described (Tenorio et al., 2018). Sections of cells in epoxy resins have high contrast and optimal morphological details. Infiltration with epoxy resin was performed at RT. All samples were polymerized at 60°C for 48h. Ultrathin sections (50-70 nm) were cut with a Leica UC6 microtome and placed on uncoated 300 mesh copper grids. Sections were contrasted with 4% uranyl acetate and Reynold’s lead citrate. Images were taken with a Tecnai G2 TEM operated at 120kV with a Ceta camera or with a Jeol 1400 operated at 120kV with a Gatan Rio camera. At least 50 cells per condition were studied by TEM.

### Immunohistochemistry

Cells were fixed with 4% paraformaldehyde and 0.1% glutaraldehyde in PBS for 1h at RT and removed from the plates in the fixative, pelleted by centrifugation and washed three times with PBS. Cell pellets were incubated 30 min with 2 % uranyl acetate in water at RT, gradually dehydrated on ice with ethanol and infiltrated at RT with LR-White acrylic resin as previously described (de Castro Martin et al., 2017). Sections of cells in acrylic resins have low contrast but an optimal preservation of protein epitopes, which is fundamental for immunolabeling studies. After polymerization at 60 degrees for 48h, the samples were sectioned using a Leica UC6 ultramicrotome. Ultrathin sections were collected on copper grids with a carbon-coated Formvar film. For immunogold labeling, unspecific binding on sections was blocked by incubation with Tris-buffered gelatin (TBG) for 5 minutes. N protein was labeled with the rabbit anti-SARS-CoV-2 nucleocapsid antibody (GXT 135357 GeneTex) diluted 1/50 in TBG for 1h. Double stranded RNA (dsRNA), an intermediate of viral replication, was labeled with the mouse monoclonal anti-dsRNA J2 antibody (10010200, English and Scientific Consulting Kft, SCICONS) diluted 1/30 in TBG for 1h. After three washes with PBS and incubation with TBG for 5 minutes, grids were labeled with a secondary antibody, corresponding to the species of the first antibody, conjugated with 10-nm colloidal gold particles (BB international) in TBG for 30 min. Sections were washed three times with PBS and five times with milliQ water and stained with saturated uranyl acetate for 20 min. After washing three times with milliQ water, grids were allowed to dry at RT. Images were taken with a Jeol 1400 TEM operated at 80kV with a Gatan One view camera or with a Jeol 1011 TEM operated at 100 kV with a Gatan ES1000W camera. At least 50 cells per condition were studied by TEM.

## RESULTS

To understand how plitidepsin exerts its mechanism of action, we first followed SARS-CoV-2 replication cycle using electron microscopy. We found no replication organelles nor viral particles in cells infected for 24 h at an MOI of 0.02, although compared to control cells, the viral infection induced some stress in cells, which showed mitochondria with swollen cristae and Golgi with swollen cisternae (not shown).

In Vero E6 cells infected for 48h at an MOI of 0.02, clusters of characteristic DMVs, which are the replication organelles of SARS-CoV-2 (Ogando et al., 2020), were present in the cytoplasm and occupied a significant area of the cytosol (**Figure 1A and B**). DMVs often had an electron lucent content with fibrillar material (**Figure 1B**). Intracellular viral particles and extracellular virions attached to the plasma membrane were abundant (**Figure 1A and C**). On average, in ultrathin sections of infected cells we found 30 intracellular viral particles and 73 extracellular particles attached to the plasma membrane per cell. Single or very few viral particles were present inside small vesicles with an electron lucent lumen while larger vacuoles with numerous viral particles were often observed (**Figure 1C**). These vacuoles that contained internal membranes and/or amorphous electron dense material might represent late endosomes/lysosomes. We seldom observed budding of viral particles into the endoplasmic reticulum (ER) (**Figure 1C**).

**Figure 1:**
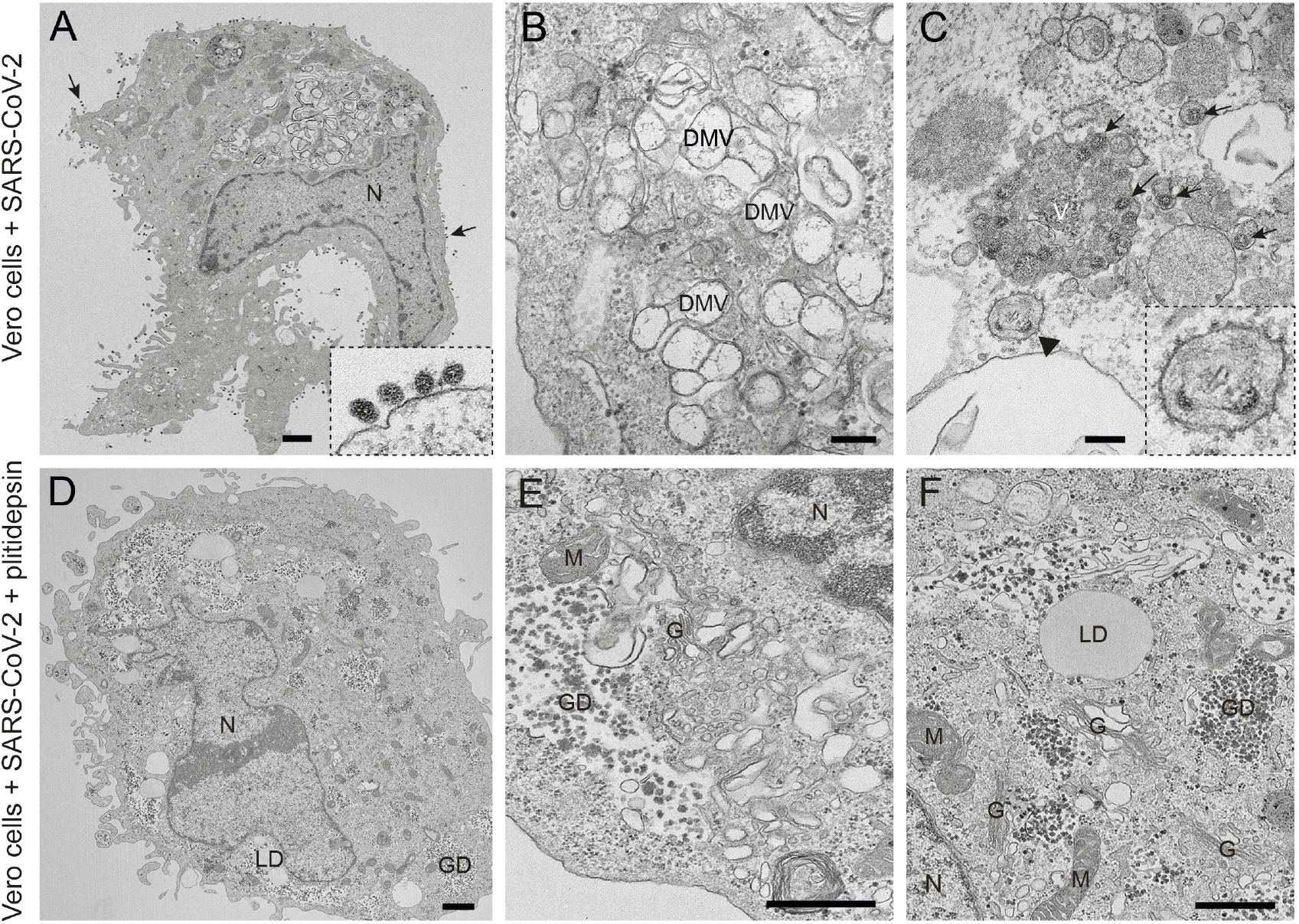
Transmission electron microscopy of Vero E6 cells infected with SARS-CoV-2 and effects of plitidepsin. (A) to (C) ultrathin sections of normally infected cells at an MOI of 0.02 and 48 h post-inoculation. (A) Cell containing a viral replication organelle made of a collection of double-membrane vesicles (DMVs) near the nucleus (N). Numerous viral particles are seen on the cell surface (arrows and inset in A). (B) DMVs have a round shape and fibrillar content. (C) Individual intracellular viral particles (arrows) are seen inside single-membrane vesicles. Large group of viruses inside a vacuole (V). Budding of viral particles in RER membranes (arrowhead and inset). (D) to (F) Ultrathin sections of Vero E6 cells infected with SARS-CoV-2 and treated with 0.2 μM plitidepsin. Low (D) and high (E and F) magnification views show that these cells contain lipid droplets (LD), large glycogen deposits (GD) and an altered Golgi complex (G) but no viral structures. M, mitochondrion. Scale bars, 1 μm in A and D; 200 nm in B, C, E and F.

When 0.2 μM of plitidepsin was added to Vero E6 cells, the drug had a profound impact on viral replication 48h later (**Figure 1D-F**). We did not observe any DMVs in the cytoplasm, nor viral particles inside cells or at the plasma membrane (**Figure 1D**). We found clusters of potential single membrane vesicles present in the cytoplasm, which are formed in early stages of SARS-CoV-2 infection (Eymieux et al., 2021a), that could however also reflect parts or a swollen Golgi (**Figure 1E**). Lipid droplets (LDs) and glycogen granules were abundant (**Figure 1F**), something also observed in mock-infected cells treated with 0.2 μM of plitidepsin (**Figure S1**). This effect was observed in cells incubated with a lower concentration of plitidepsin, that is 0.05 μM, only less pronounced (**Figure S2**).

We then studied in detail the effects of 0.05 or 0.2 μM of plitidepsin on the assembly of SARS-CoV-2 replication organelles or DMVs. Of note, these two concentrations are close to the IC_50_ and IC_90_ estimated for SARS-CoV-2 induced cytopathic effect on Vero E6 cells (Rodon et al., 2021). When plitidepsin was added to Vero E6 cells at the same time of viral infection (MOI of 0.02 for 48h), both concentrations had a deep impact on the biogenesis of DMVs (**Figure 2**). Cells infected without plitidepsin contained numerous, typical DMVs with fibrillar content surrounded by viral particles inside single-membrane vesicles (**Figure 2A-C**) but when the drug was added no DMVs were found (**Figure 2D-I**). Already at the low concentration, no DMVs were found in the cytosol (**Figure 2D-F**). Single membrane vesicles were present but they did not contain fibrillar content in their lumen as opposed to infected cells, and this could reflect a part of a swollen Golgi or ER (**Figure 2F and I**). These morphological analyses on plitidepsin-treated cells failed to identify any structures implicated in SARS-CoV-2 replication that were unambiguously detected in untreated infected cells (**Figures 1A-C and 2A-C**).

**Figure 2:**
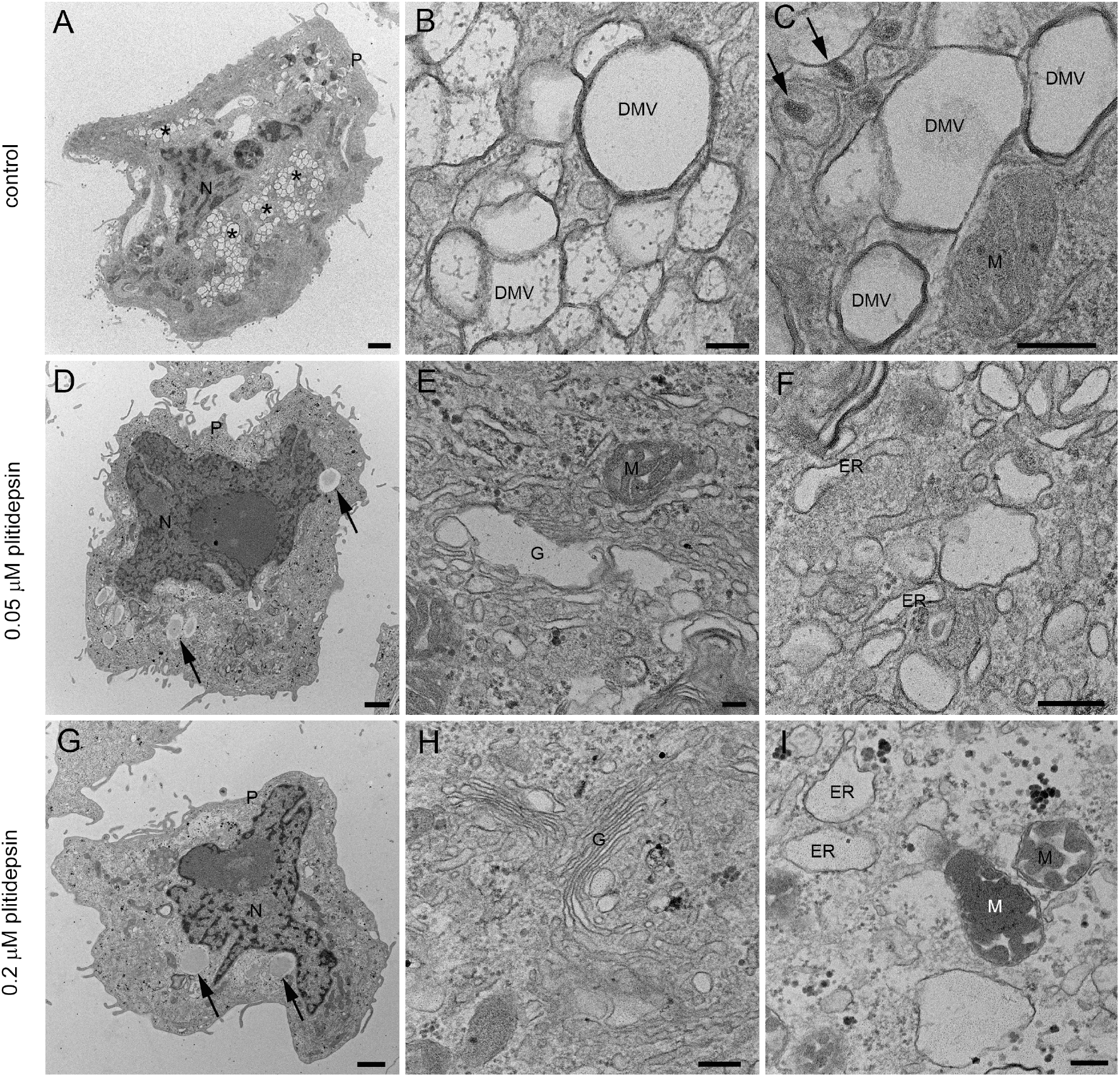
Transmission electron microscopy of Vero E6 cells infected with SARS-CoV-2 in the absence or presence of two doses of plitidepsin. (A) to (C) Normally infected cells. Low (A) and high (B and C) magnification of ultrathin sections of cells infected 48 h at an MOI of 0.02 (A) Groups of characteristic double membrane vesicles (DMVs; asterisks) occupy large areas of infected cells. (B) Typical DMVs exhibit electron dense membranes and sometimes a fibrillar content. (C) Single membrane vesicles with viruses (arrows) are often found in the close vicinity of DMVs. (D) to (F) Low (D) and high (E) and (F) magnification of Vero E6 cells infected with SARS-CoV-2 48 h at an MOI of 0.02 and treated with 0.05 μM plitidepsin. Cells contain lipid droplets (arrows in D), swollen Golgi stacks (G), mitochondria with swollen cristae (M) and swollen endoplasmic reticulum (ER) cisternae. No viral structures are seen. (G) to (I) Cells infected with SARS-CoV-2 48 h at an MOI of 0.02 and treated with 0.2 μM plitidepsin. Cells contain lipid droplets (arrows in G), glycogen deposits (GD in H and I), swollen ER and altered mitochondria (I). N, nucleus; P, plasma membrane. Scale bars, 1 μm in A, D and G; 200 nm in B, C, E, F, H and I.

The absence of structures implied in viral replication was confirmed by immunodetection of two key products of SARS-CoV-2: the nucleocapsid and double stranded RNA (dsRNA). The nucleocapsid participates in replication and translation of viral RNA and maintains the structure of ribonucleoprotein complex, whereas the dsRNA is synthesized by the viral polymerase in the host cell, once viral replication has started in DMVs. For both targets, no signal by immunogold labeling was detected in non-infected cells, either non treated with plitidepsin or incubated with low or high concentration of plitidepsin (**Figure S3 and S4**). However, virus-infected cells showed a strong labeling for the nucleocapsid in the cytosol and in viral particles that accumulated intra-as well as extracellularly (**Figure 3A-C**). Coherent with the morphological observations, after application of plitidepsin at 0.05 μM, no signal was detected by antibody labelling for nucleocapsid (**Figure 3D-F**). The absence of labelling was also found for the high concentration of plitidepsin (0.2 μM) (**Figure 3G-I**). In SARS-CoV-2 infected cells without plitidepsin, the labeling for dsRNA was restricted to the lumen of large electron lucent vesicles, which represent the DMVs (**Figure 4A-C**). Due to the low contrast of ultrathin sections of cells embedded in the LR-White acrylic resin, the fibrillar content in the lumen of DMVs is not visible in these samples but the labeling of dsRNA marked them unequivocally as DMVs. After application of Plitidepsin at 0.05 or 0.2 μM, no signal for dsRNA was detected by immunogold labelling (**Figure 4D-I**). Overall, immunodetection of viral proteins and genetic material confirmed that the morphological analysis performed corresponded to viral replication structures and that plitidepsin completely blocked the assembly of the viral replication organelles or DMVs and the accumulation or viral nucleocapsid protein and dsRNA. Taken together these results provide evidence that plitidepsin inhibits intracellular viral replication and abrogates the formation of DMVs.

**Figure 3:**
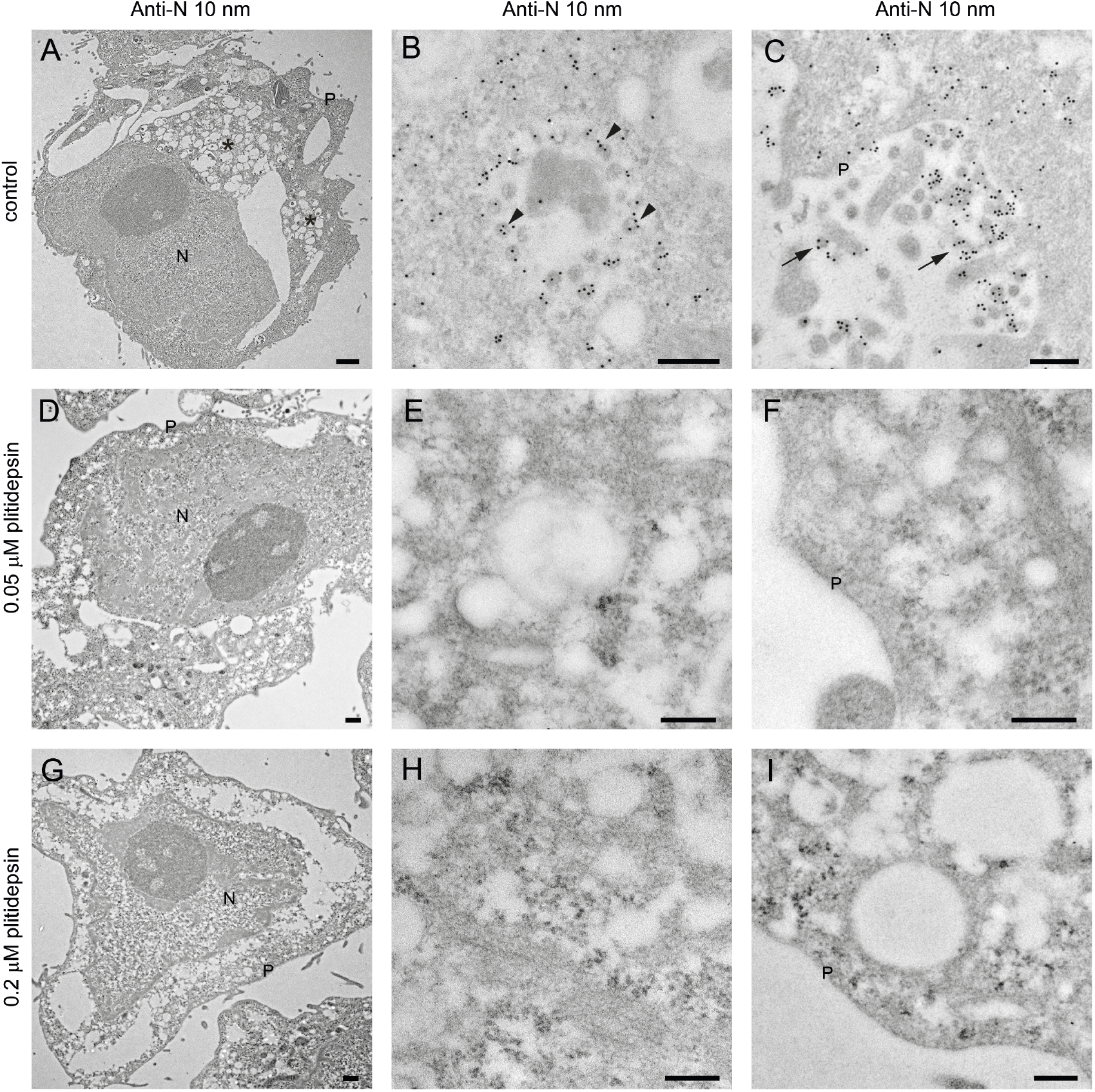
Immunogold detection of SARS-CoV-2 nucleocapsid protein in infected cells in the absence or presence of two doses of plitidepsin. Ultrathin sections of cells were incubated with anti-N primary antibody followed by a secondary antibody conjugated with 10 nm colloidal gold particles and visualized by TEM. (A) to (C) Ultrathin sections of cells infected for 48 h with SARS-CoV-2 at an MOI of 0.02. (A) Low magnification image of an infected cell with characteristic DMVs (asterisks) near the nucleus (N). (B) High magnification image of a group of intracellular viruses (arrowheads) inside a vacuole (V). Viral particles and areas of cytosol are labelled. (C) Labelled extracellular virions (arrows) on the cell surface. (D) to (F) Cells infected 48h with SARS-CoV-2 at an MOI of 0.02 and treated with 0.05 μM plitidepsin. Low (D) and high (E) and (F) magnification images of cells show no labeling. P, plasma membrane. (G) to (I) Cells infected 48 h with SARS-CoV-2 at an MOI of 0.02 and treated with 0.2 μM plitidepsin. Low (G) and high (H) and (I) magnification images of cells show no labeling. Scale bars, Scale bars, 0.5 μm in A, D and G; 200 nm in B, C, E, F, H and I.

**Figure 4.**
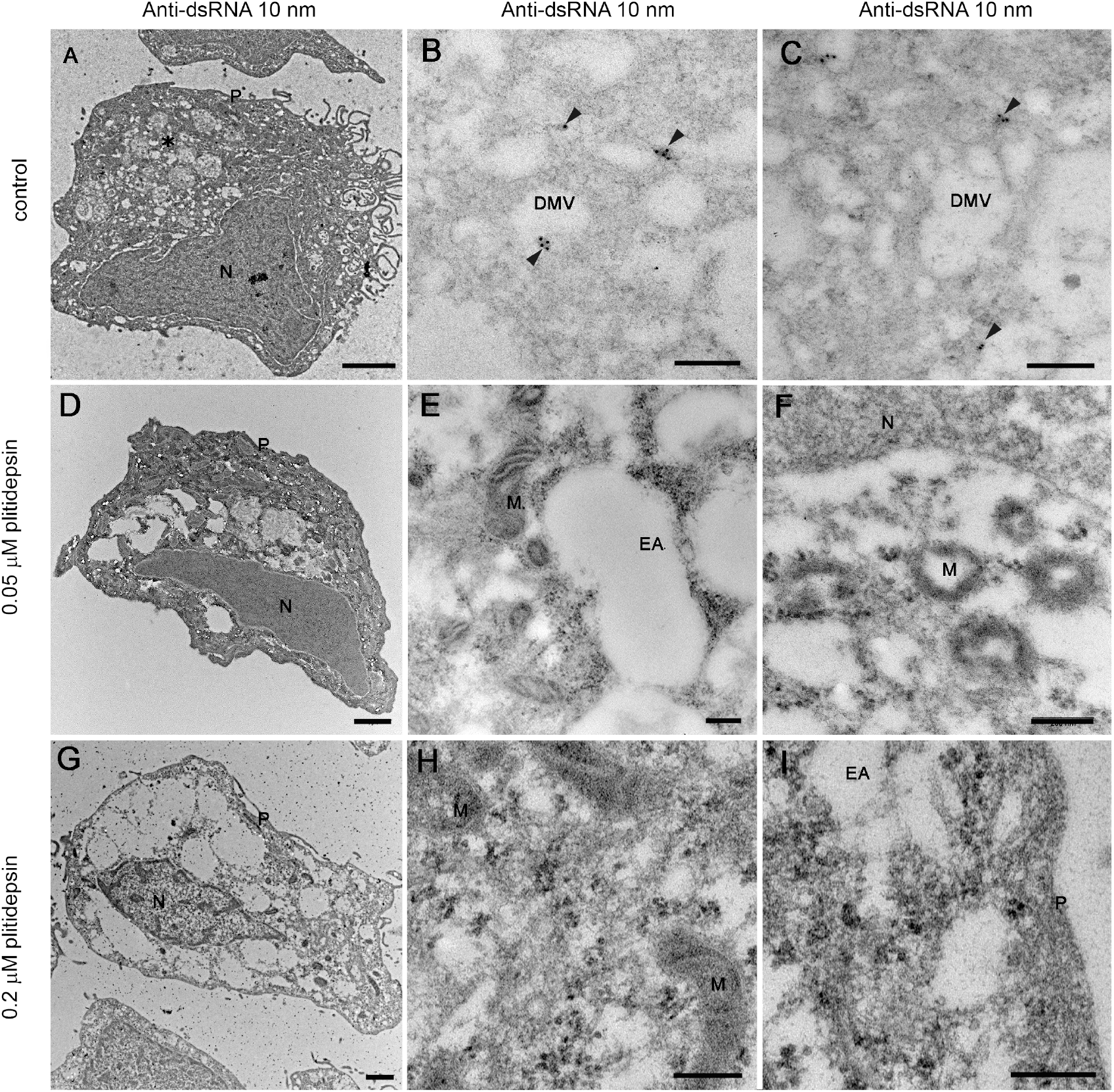
Immunogold labelling of SARS-CoV-2 dsRNA in infected cells in the absence or presence of two doses of plitidepsin. Ultrathin sections of cells were incubated with anti-dsRNA primary antibody followed by a secondary antibody conjugated with 10 nm colloidal gold particles and visualized by TEM, (A) to (C) Ultrathin sections of cells infected 48 h with SARS-CoV-2 at an MOI of 0.02. (A) Low magnification image of an infected cell with characteristic DMVs (asterisk) near the nucleus (N). (B) and (C) High magnification images of labelled DMVs (arrowheads). (D) to (F) Cells infected 48 h with SARS-CoV-2 at an MOI of 0.02 and treated with 0.05 μM plitidepsin. Low (D) and high (E) and (F) magnification images of cells show no labeling. to (I) Cells infected 48 h with SARS-CoV-2 at an MOI of 0.02 and treated with 0.2 μM plitidepsin. Low (G) and high (H) and (I) magnification images of cells show no labeling. Scale bars, Scale bars, 1 μm in A, D and G; 200 nm in B, C, E, F, H and I.

## DISCUSSION

The TEM analysis of SARS-CoV-2-infected Vero E6 cells performed herein reveals the formation of DMV structures characteristic of the viral replication organelle, and the accumulation of intracellular viral particles in small vesicles and large vacuoles as well as extracellular virions at the plasma membrane. These structures have been described before as characteristic features of SARS-CoV-2 infection in cell culture (Eymieux et al., 2021a, 2021b; Ogando et al., 2020). In this study, infected cells displayed a clear specific labelling for both the nucleocapsid viral protein and the dsRNA viral replication intermediate. Specific labelling of two key viral products is highly relevant in a context in which TEM analysis of SARS-CoV-2-infected samples has led to the misidentification of viral particles and to ambiguous results (Dittmayer et al., 2020; Miller and Goldsmith, 2020). Together, the morphological analysis along with the immunogold detection of two complementary SARS-CoV-2 products performed herein provided conclusive evidence that the subcellular structures detected in infected cells represented viral replication organelles and virus progeny. In the present set of experiments, plitidepsin abrogated the formation of DMVs and the detection of nucleocapsid and dsRNA viral products in SARS-CoV-2-infected Vero E6 cells. Since plitidepsin interferes with the host factor eEF1A implicated in RNA translation (Losada et al., 2016), it is worthwhile to know the impact of this mechanism on the formation of DMVs.

The antiviral effect of plitidepsin against SARS-CoV-2 is mediated throughout inhibition of the host protein eEF1A. siRNA silencing of eEFA1A in host cells induces a significant reduction in the nucleocapsid protein levels, as well as a reduction in the viral RNA, which demonstrates a direct involvement of eEF1A in the viral replication (Zhang et al., 2014). In addition, the exposure of plitidepsin to cells at the moment of SARS-CoV-2 infection reduced the viral replication and transcription, which induced a reduction in the accumulation of genomic RNA and sub-genomic N RNA viral levels, respectively (White et al., 2021). Due to the known high sensitivity of coronavirus to translation inhibitors (Bojkova et al., 2020; van den Worm et al., 2011), especially at an early stage of infection, a plitidepsin-mediated inhibition of SARS-CoV-2 protein translation process could represent one possible mechanism, which could lead to the abrogation of DMV formation. The results gathered from the present experiments demonstrate that a concentration as low as 50 nM of plitidepsin, completely abolished the formation of DMVs, and the synthesis of N protein and viral dsRNA in SARS-CoV-2 infected Vero E6 cells both at 24 and 48 h post-inoculation. The plitidepsin-mediated anti-SARS-CoV-2 activity at such a low concentration may be the result of a highly specific inhibition of genomic RNA viral translation. This specific antiviral effect is explained by the lack of plitidepsin-mediated protein synthesis inhibition in eukaryotic cells at high concentrations (e.g., 450 nM or higher), far above those used herein (Losada et al., 2016). Inhibition of the viral protein translation may affect several essential proteins, such as non-structural proteins, whose predicted transmembrane domains are key for DMV formation during the replication of distinct coronaviruses (Angelini et al., 2013; Oudshoorn et al., 2017; Wolff et al., 2020a). It is tempting to speculate that the lack of translation of these viral proteins could directly abrogate the formation of these replication–transcription complexes necessary for DMV formation. This possible mechanism is supported by prior observations where plitidepsin treatment decreased nucleocapsid content and reduced subgenomic RNA detection (Rodon et al., 2021; White et al., 2021). An alternative mechanism has been recently proposed with eEF1A binding to membranes and thereby recruiting a subset of proteins related to DMV formation (Carriles et al., 2021).

In the present study, TEM morphological analysis coupled to immunogold labeling of SARS-CoV-2 products have demonstrated to be a valuable tool to better understand the antiviral effects, as previously reported for other viruses (García-Serradilla, 2021; Sachse et al., 2019). Remarkably, the plitidepsin-induced complete blockade of the assembly of viral structures detected here by electron microscopy is rather unique. In the presence of non-toxic, high concentrations of other antiviral drugs, SARS-CoV-2 often assembles small amounts of DMVs and viral particles (Izquierdo-Useros, Cerón and Risco, unpublished results). The generated insights warrant new experiments that aim to better understand the mechanism of plitidepsin-induced antiviral activity. This knowledge will be crucial to identify the mechanism of action for promising compounds that interfere with host factors whose implication in key biological processes can be applied as a pan-antiviral strategies.

## FINANCIAL SUPPORT

This research was funded by Pharma Mar, which commercializes Aplidin/Plitidepsin. The authors also acknowledge the crowdfunding initiative #Yomecorono (https://www.yomecorono.com). N.I-U. is supported by grant PID2020-117145RB-I00 from the Spanish Ministry of Science and Innovation. C.R. is supported by grant RTI2018-094445-B-I00 (MCI/AEI/FEDER, UE) from the Spanish Ministry of Science and Innovation.

## ACKNOWLEDGEMENTS

We acknowledge J. Pedroza from the CMCiB for his constant help at the BSL3 facility.

## COMPETING INTEREST

P.A., J.V-A and N.I-U. are inventors in a patent application related to Aplidin/Plitidepsin (EP20382821.5). Unrelated to the submitted work, N.I-U. reports institutional grants from Grifols, Dentaid, Hipra and Palobiofarma. The authors declare that no other competing financial interests exist.

## DATA AVAILABILITY

Data related to this work is available from corresponding authors upon reasonable request.

## Supplementary Materials

**Supplementary Figure 1.** Effects of plitidepsin in the morphology of mock-infected Vero E6 cells. (A) and (B) Low and high magnification images of control Vero E6 cells with normal nucleus (N), Golgi complex (G) and mitochondria (M). (C) and (D) Low and high magnification images, respectively, of Vero E6 cells treated 48 h with 0.2 μM plitidepsin. Cells show lipid droplets (LD), altered mitochondria (M), glycogen deposits (GD) and swollen Golgi (G). Scale bars, 500 nm.

**Supplementary Figure 2.** Dose-dependent effects of plitidepsin in mock-infected Vero E6 cells. (A) to (C) control cells. Low (A) and high (B) and (D) magnification images, respectively, of untreated cells. (D) to (F) Vero E6 cells incubated 48 h with 0.05 μM plitidepsin. Low (D) and high (E) and (F) magnification images, respectively. Arrows in (D) point to lipid droplets. (G) to (I) Vero E6 cells incubated 48 h with 0.2 μM plitidepsin. Low (G) and high (H) and (I) magnification images, respectively. N, nucleus; P, plasma membrane; G, Golgi complex; ER, endoplasmic reticulum; LB, lamellar body; GD, glycogen deposits; LD, lipid droplet. Scale bars, 1 μm in A, D and G; 200 nm in B, C, E, F, H and I.

**Supplementary Figure 3.** Immunogold cytochemical control of anti-nucleocapsid (N) antibody in non-infected Vero E6 cells, with and without plitidepsin. Ultrathin sections of cells were incubated with anti-N primary antibody followed by a secondary antibody conjugated with 10 nm colloidal gold particles and visualized by TEM. (A) to (C) control, non-infected Vero E6 cells in the absence of plitidepsin. Low (A) and high (B) and (C) maginification images. (D) to (F) Vero E6 cells treated 48 h with 0.05 μM plitidepsin. Low (D) and high (E) and (F) magnification images. (G) to (I) Vero E6 cells treated 48 h with 0.2 μM plitidepsin. Low (G) and high (H) and (I) magnification images. Cells present no unspecific background. N, nucleus; P, plasma membrane; M, mitochondrion; LB, lamellar body. Scale bars, 0.5 μm in A, D and G; 200 nm in B, C, E, F, H and I.

**Supplementary Figure 4.** Immunogold cytochemical control of anti-dsRNA antibody in non-infected Vero E6 cells, with and without plitidepsin. Ultrathin sections of cells were incubated with anti-dsRNA primary antibody followed by a secondary antibody conjugated with 10 nm colloidal gold particles and visualized by TEM. Low (A) and high (B) and (C) magnification images. (D) to (F) Vero E6 cells treated 48 h with 0.05 μM plitidepsin. Low (D) and high (E) and (F) magnification images. (G) to (I) Vero E6 cells treated 48 h with 0.2 μM plitidepsin. Low (G) and high (H) and (I) magnification images. Cells present no unspecific background. N, nucleus; P, plasma membrane; M, mitochondrion. Scale bars, 1 μm in A, D and G; 200 nm in B, C, E, F, H and I.

