## Supplementary figures and images for "Unraveling the antiviral activity of plitidepsin by subcellular and morphological analysis"

### Supplementary Figures

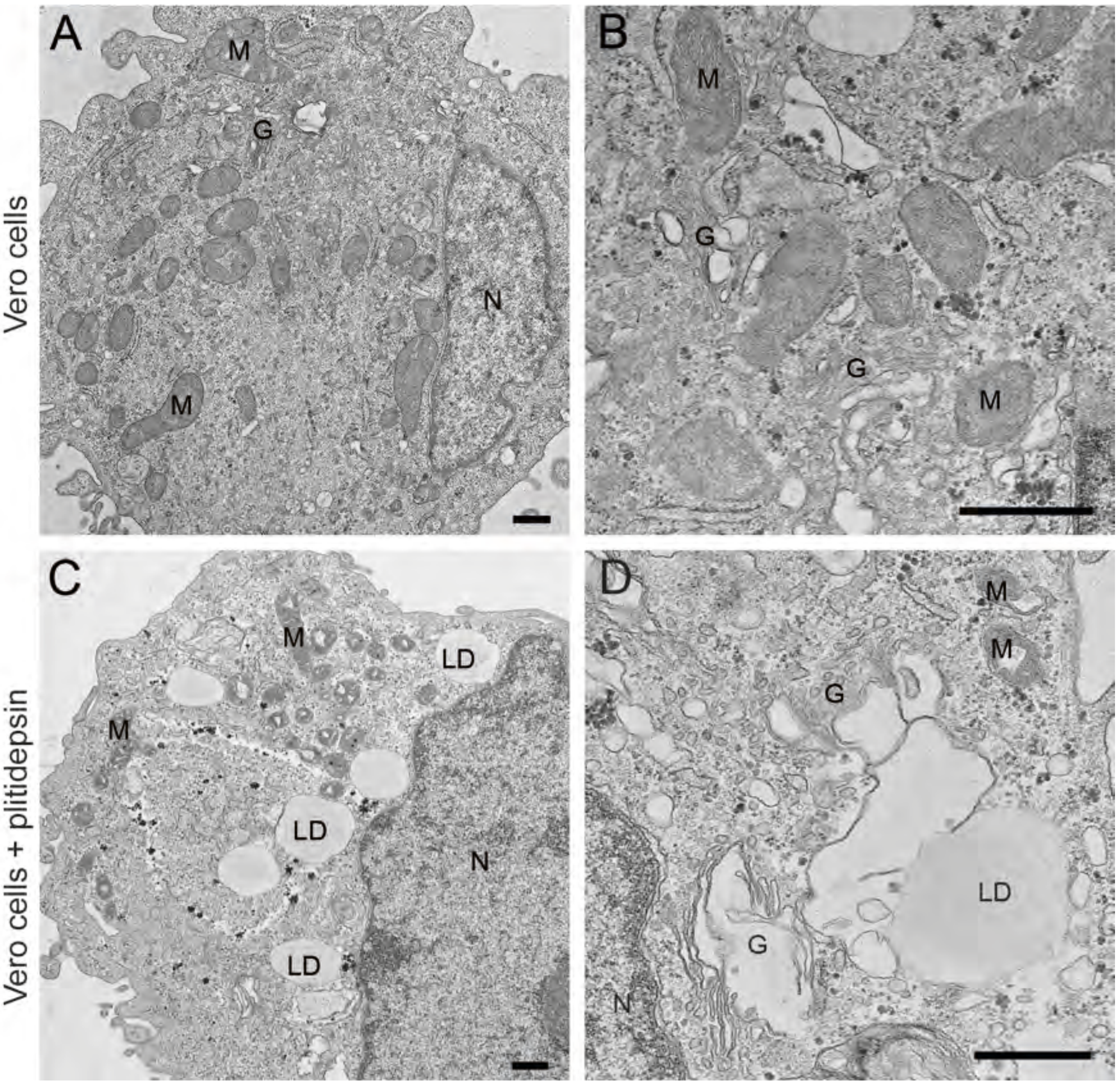

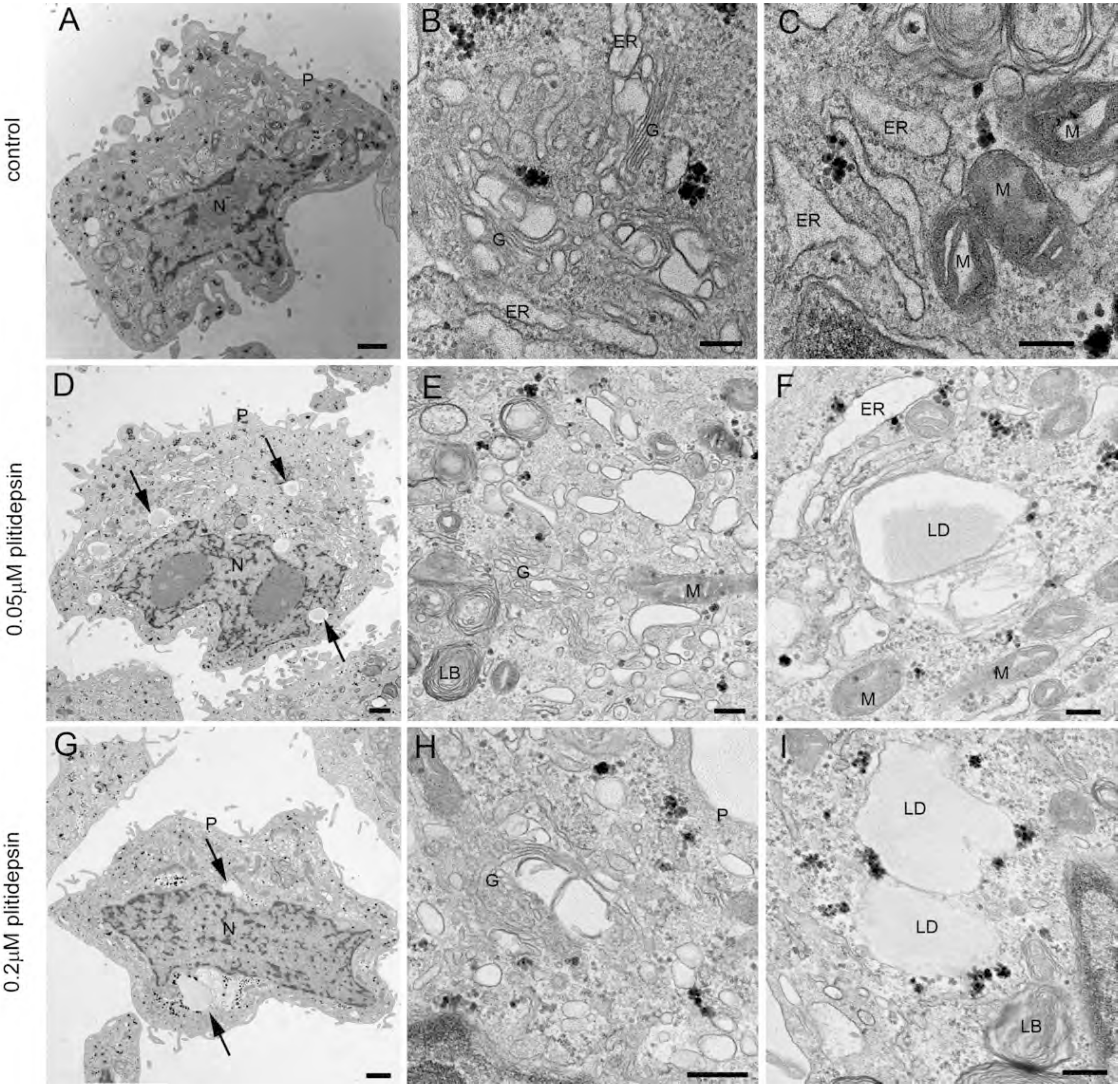

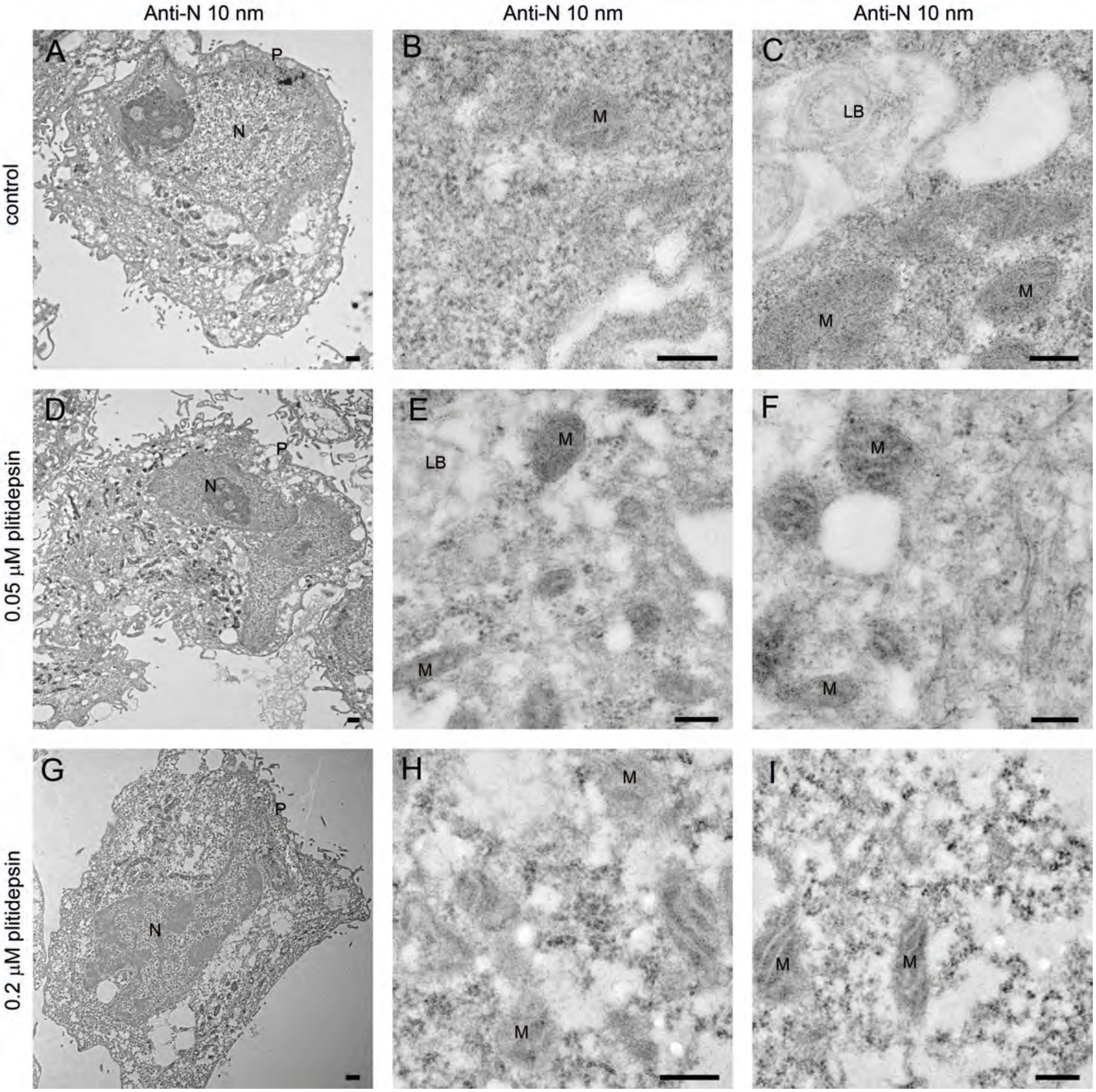

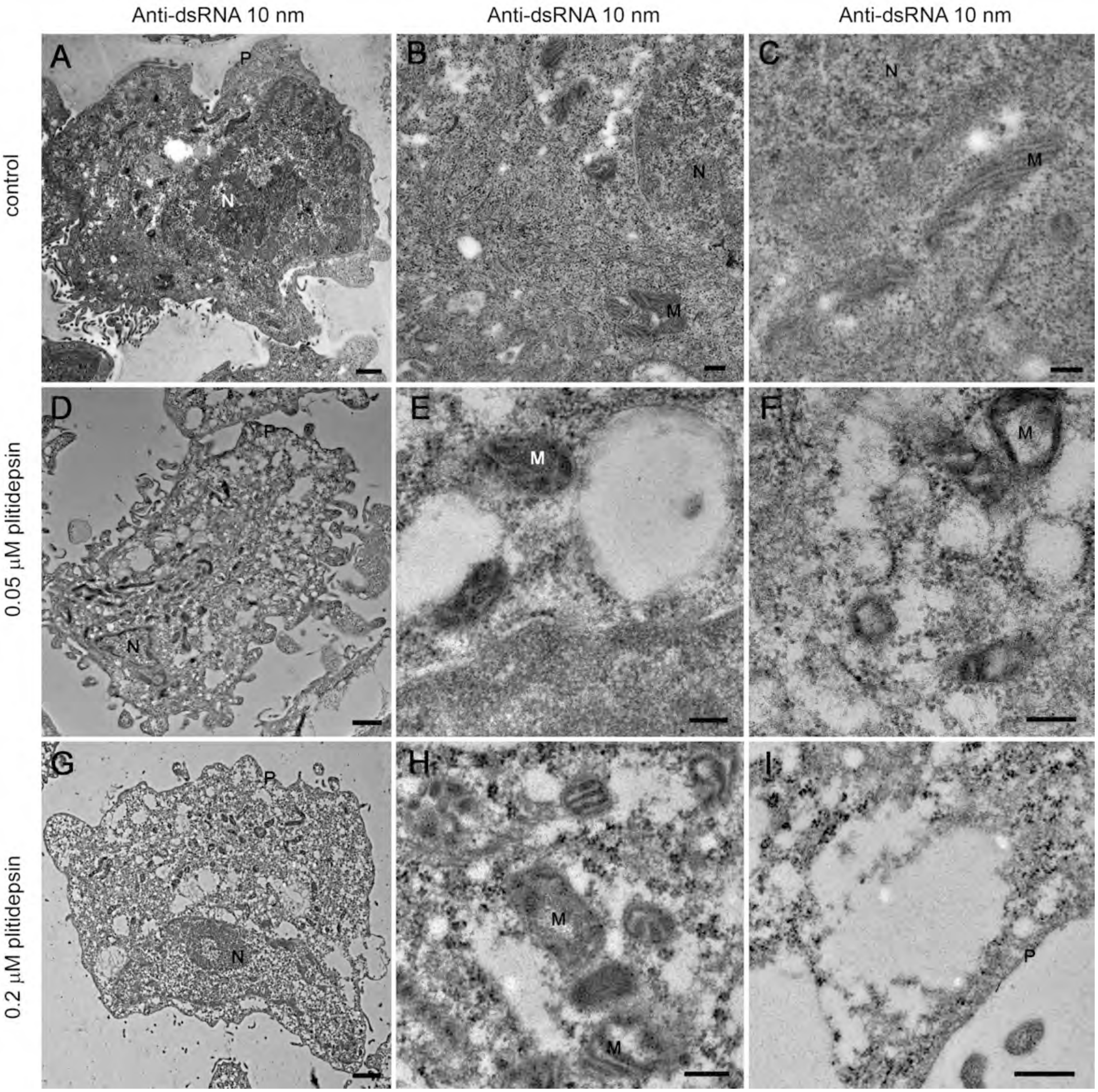
